# Physical epistatic landscape of antibody binding affinity

**DOI:** 10.1101/232645

**Authors:** Rhys M. Adams, Justin B. Kinney, Aleksandra M. Walczak, Thierry Mora

## Abstract

Affinity maturation produces antibodies that bind antigens with high specificity by accumulating mutations in the antibody sequence. Mapping out the antibody-antigen affinity landscape can give us insight into the accessible paths during this rapid evolutionary process. By developing a carefully controlled null model for noninteracting mutations, we characterized epistasis in affinity measurements of a large library of antibody variants obtained by Tite-Seq, a recently introduced Deep Mutational Scan method yielding physical values of the binding constant. We show that representing affinity as the binding free energy minimizes epistasis. Yet, we find that epistatically interacting sites contribute substantially to binding. In addition to negative epistasis, we report a large amount of beneficial epistasis, enlarging the space of high-affinity antibodies as well as their mutational accessibility. These properties suggest that the degeneracy of antibody sequences that can bind a given antigen is enhanced by epistasis — an important property for vaccine design.

---

To ensure a reliable response and to neutralize foreign pathogens, the adaptive immune system relies on affinity maturation. In this process, antibody receptors expressed by B cells undergo an accelerated Darwinian evolution through random mutations and selection for affinity against foreign epitopes [1]. Mature antibodies can accumulate up to 20% hypermutations, leading to up to a 10,000 fold improvement in binding affinity [2]. Affinity maturation also produces broadly neutralizing antibodies that target conserved regions of the pathogen, of particular importance for vaccine design against fast evolving viruses [3]. Despite extensive experimental and theoretical work, the key determinants of antibody specificity and evolvability are still poorly understood, mainly because the sequence-to-affinity relationship is difficult to measure comprehensively or to predict computationally [4].

A major confounding factor in characterizing the sequence dependence of any protein function, including affinity, is the pervasiveness of epistasis, the phenomenon by which different mutations interact with each other [5]. Theory [6-8] and genomic data [9] suggest that inter- and intragenic epistasis plays a major role in molecular evolution, by constraining the set of accessible evolutionary trajectories towards adapted phenotypes [10–14], enhancing evolvability through stabilizing mutations [15, 16], or slowing down adaptation by the law of diminishing returns [17, 18]. Evidence for epistasis in antibody affinity include direct observations of cooperativity between mutations [19, 20], the dependence of mutational effects on sequence background [21], and statistical covariation of residues in large sequence datasets [22, 23].

Intragenic epistasis has mostly been studied either by measuring the fitness of all possible mutational intermediates between two variants [10, 24-27], or by comparing the effect of mutations in different backgrounds [21, 28, 29]. Many such studies rely on a particular measure of fitness rather than a well-defined physical phenotype. Deep mutational scans (DMS) [30] can comprehensively map out the epistatic landscape of many genetic variants [14, 31, 32]. However, most DMS methods rely on noisy selection and do not measure the biophysical quantity of interest directly [33], introducing both nonlinearities and noise that could be misinterpreted as epistasis.

Here we analyze the detailed epistatic landscape of an antibody’s binding free energy to its cognate antigen (the 4-4-20 antibody fragment against fluorescein), using data previously obtained by Tite-Seq, a recently introduced DMS variant that accurately measures protein binding affinity in physical units of molarity [34]. By comparing to a simple additive model of mutations on the binding free energy, and carefully controling for measurement noise and nonlinearities, we find that epistasis significantly contributes to the antibody’s affinity. This epistasis is not uniformly distributed, but instead favors certain residue pairs across the protein. We use our results to analyze how epistasis both constrains and enlarges the set of possible evolutionary paths leading to high-affinity sequences.

## RESULTS

### Position Weight Matrix model of affinity

We analyzed data from [34] (https://github.com/jbkinney/16_titeseq), where Tite-Seq was applied to measure the binding affinities of variants of the 4-4-20 fluorescein-binding scFv antibody, henceforth called ‘wildtype’. Libraries were generated by introducing mu-tations to either the CDR1H or CDR3H domains re-stricted to 10 amino acid stretches called 1H and 3H (Fig. 1A). All single amino acid mutants, 1100 random double amino acid mutants, and 150 triple amino acid mutants were generated in multiple synonymous variants and measured, (Fig. 1B). Using a combination of yeast display and high-throughput sequencing at various antigen concentrations, Tite-Seq yielded the binding dissociation constant *K_D_* (in M or mol/L) of each variant with the fluorescein antigen.

**Fig. 1.**
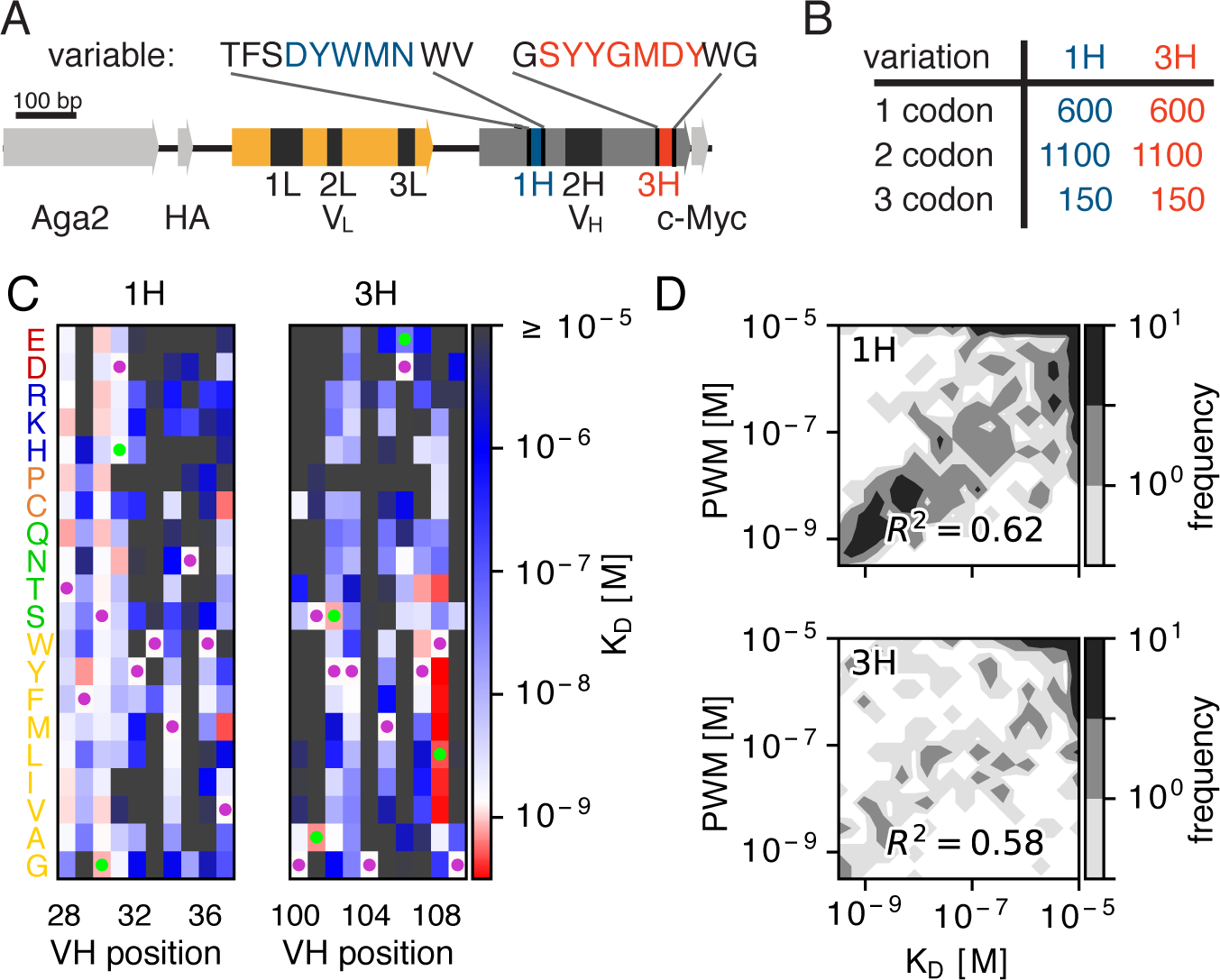
Additive model of binding affinity. **(A)** 4-4-20 scFv antibody sequence. Six complementarity determining regions (CDR: 1L, 2L, 3L, 1H, 2H, 3H) are particularly important for antibody binding affinity. **(B)** A library of antibody sequences with mutations in 10 amino-acid regions around the CDR1H and CDR3H domains were expressed using yeast display. Using Tite-Seq, the binding constants *K_D_* of all 600 single codon mutants, 1100 random double codon mutants, and 150 random triple codon mutants, were measured. **(C)** The *K_D_* of single mutants for 1H and 3H domains were used to create position weight matrices (PWM) to predict the affinity of double and triple mutants. **(D)** Comparison between the PWM prediction and the measurement of *K_D_* on double and triple mutants. The PWMs explained a significant portion of the variance, as quantified by the explained variance *R*^2^ (*p* < 10^−61^ for CDR1H, *p* < 10^−48^ for CDR3H, F-test for reduction in variance due to PWM). PWMs trained from the binding free energy, *F* = ln(*K_D_*/*c*_0_), outperformed PWM trained from *K_D_* (Fig. S1). We looked for the optimal nonlinear transformation of *K_D_* maximizing the PWM fits (Methods and Fig. S2 for validation on simulated data) and found that PWMs perform almost ideally when trained from the binding free energy (see Fig. S3).

We first tried to predict the *K_D_* of double and triple mutants from single mutant measurements. Mutations are expected to act on the binding free energy in an approximately additive way [32, 35]. One may thus write the free energy of binding, *F* = ln(*K_D_/c*_0_) (defined up to constant in units of *k_B_T*), as a sum over mutations in the mutagenized region, s = (*s*_1_,…, *s_l_*:

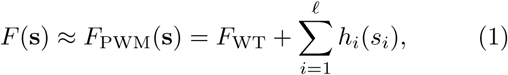

where *F_WT_* is the wildtype sequence energy, and *h_j_*(*s_j_*) is the effect of a mutation at position *i* to residue *S_i_*. The elements of the Position-Weight Matrix (PWM) *h_i_*(*s*) are obtained from the *K_D_* of single mutants shown in Fig. 1C. Since Tite-Seq measurements are limited to values of *K_D_* ranging from 10^−95^ to 10^−5^, for consistency PWM predictions outside this range were set to the boundary values. The PWM was a fair predictor of double and triple mutants (Fig 1D), accounting for 62% (*p* < 10^−61^, F-test) of the variance for 1H mutants and 58% (*p* < 10^−48^, F-test) of the variance of 3H mutants.

The unexplained variance missed by the PWM model may have three origins: measurement noise, epistasis, or nonlinear effects. The last case corresponds to the hypothesis of additivity not being valid for *F* = ln(*K_D_/c*_0_), but for some other nonlinear transformation of *F*. Such a nonlinearity, also called “scale,” can lead to spurious epistasis [5, 36]. We first checked that additivity did not apply to the untransformed dissociation constant, *K_D_*: a PWM model learned from *K_D_* instead of *F* could only explain 34% of the variance of all 1H and 3H multiple mutants, down from 62% when learning from *F* (Fig. S1). We then looked for the non-linear transformation *E*(*F*) that would give the PWM model with the best predictive power (Methods and Fig. S2). This optimization yielded only a modest improvement to 65% of the explained variance. In addition, the optimal function *E* was very close to the logarithm (*R*^2^ = 97%, Fig. S3). Since nonlinear effects do not play a significant role, henceforth we only consider the PWM model defined on the free energy.

### Epistasis affects affinity

To identify epistasis, we estimated the difference between the measured binding free energies of double and triple mutants, *F*(s), and the PWM prediction, *F*_PWM_(s). However, these small differences can be confounded by measurement noise, which can be mistaken for epistasis. To control for this noise, we defined Z-scores between two estimates of the free energy, *F_a_* and *F_b_,* as 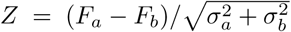, where 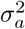 and 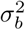 are their estimates of uncertainty. We first computed Z-scores between independent estimates of the same free energy using synonymous variants (*Z*_error_, Methods). We found that the distribution of *Z*_error_ was normal with variance ≈ 1 (Fig. 2A, orange line), as expected from Gaussian measurement noise.

**Fig. 2.**
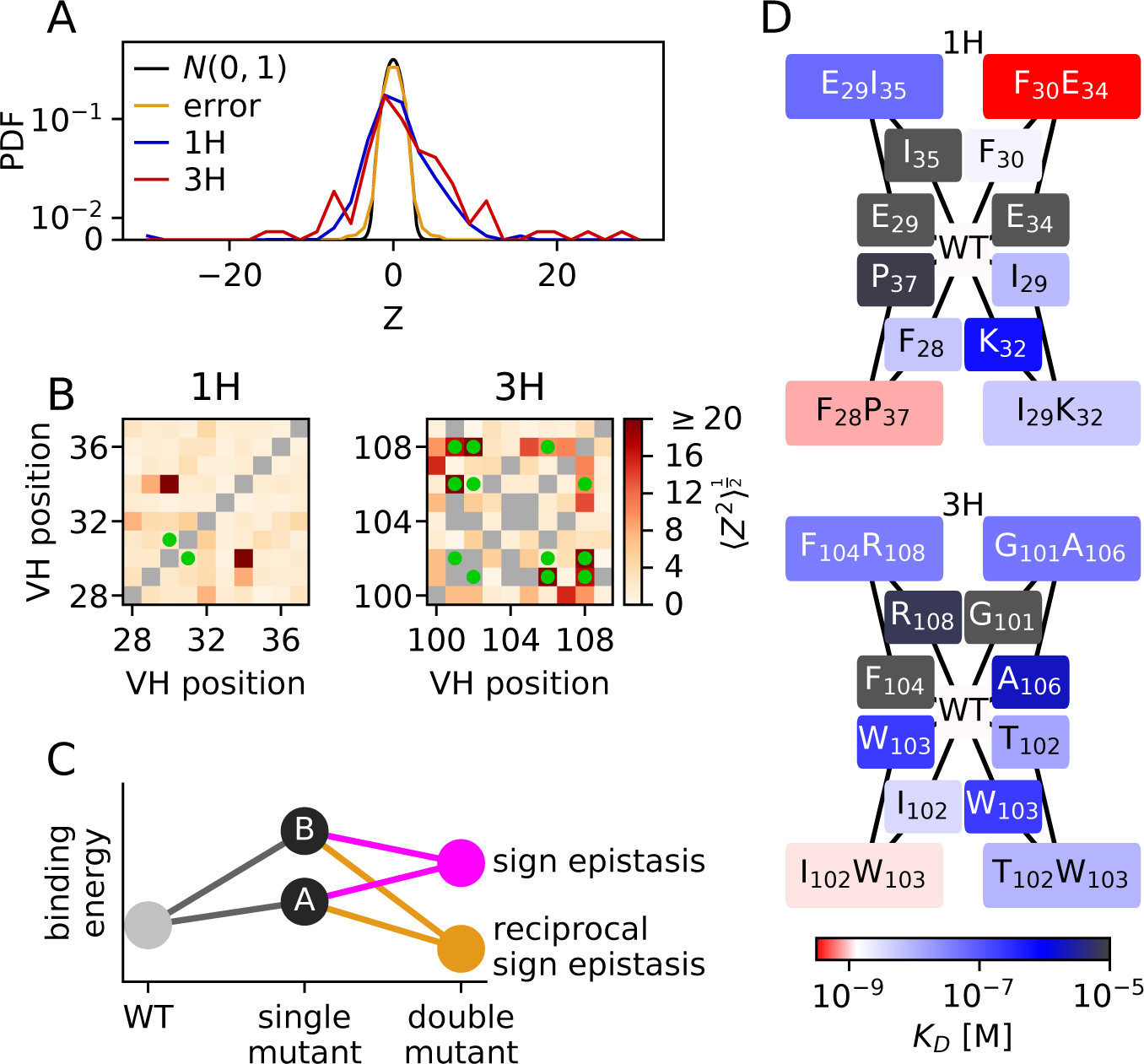
Quantification of epistasis. Epistasis is defined as deviation from the PWM model, which assumes an additive effect of single mutations on the binding free energy *F* = ln(*K_D_/c*_0_) expressed in units of *k_B_T*. **(A)** Distribution of Z-scores, defined as the normalized deviation from the PWM prediction, 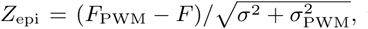 where *σ*^2^ and 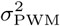 are the estimated errors on *F* and *F*_PWM_. Positive Z-scores indicate epistasis increased affinity. The Z score standard deviation was much higher than expected from measurement errors (*Z*_error_) for CDR1H (3.34, *p* < 10^−33^, Levene’s test) and CDR3H (5.44, *p* < 10^−52^), meaning that the discrepancy between the PWM and measurement is mainly due to true epistasis. **(B)** Standard Z-score deviation for each pair of positions along the sequence. This deviation is higher at pairs of positions mutated in the super-optimized 4m5.3 antibody (green dots) in 3H (*p* = 0.005, Mann-Whitney), but not in 1H (*p* = 0.23). Pairs of positions with large epistatic effects are shown on the wild-type crystal structure in Fig. S5. There is a weak correlation between epistasis and distance between the residues (Fig. S6). **(C)** Schematic representation of sign and reciprocal sign epistasis for a beneficial interaction. **(D)** Representative examples of sign epistasis as identified by Z-score. All 44 examples of beneficial sign epistasis with double mutant *K_D_* ≤ 10^−6^ may be found in `S1_table_sign_epistasis.csv`, and summarized in tables S1 and S2. Examples of deleterious sign epistasis are shown in figure S4.

We then estimated the effect of epistasis by calculating Z-scores (*Z*_epi_) from the difference between the PWM prediction, *F*_PWM_ (Eq. 1), and the measured *F*. The resulting distributions of Z-scores (Fig. 2A, blue and red lines) had much larger variances than expected from measurement noise (standard deviation 3.34 for 1H, and 5.44 for 3H), indicating strong epistasis. These epistatic effects were on average slightly beneficial (positive *Z*): 25% of double mutants inside the reliable readout boundaries (10^−9.5^*M* ≤ *K_D_* ≤ 10^−5^*M*) showed significant beneficial epistasis (*Z*_epi_ > 1.64, *p* = 0.05), and 20% significant deleterious epistasis (*Z*_epi_ < −1.64). Comparing the variance of *Z*_epi_ with that of *Z*_error_ gives a large fraction of the unexplained variance that is attributable to epistasis, 1 — Var(*Z*_error_)/Var(*Z*_epi_) = 89% for 1H, and 96% for 3H.

To determine whether certain positions along the sequence concentrated epistatic effects, we computed the mean squared Z-score for all double mutations at each pair of positions (excluding median boundary values), revealing a complex and heterogeneous landscape of epistasis (Fig. 2B and Fig. S5 for the epistasis magnitude superimposed on the wildtype’s crystal structure). CDR3H, which interacts directly with the antigen, is observed to have more epistatically interacting sites than CDR1H. Interestingly, the three most epistatic pairs in 3H — between positions 101, 106 and 108 — are mutated in the previously described super-optimized 4m5.3 antibody [37] (mutations shown in green in Fig. 1B), consistent with previous suggestions that positions 101 and 106 interact together and with position 108 via hydrogen bonds [19, 34]. Epistasis is usually expected between residues that are in contact in the protein structure [38-42], as for instance between positions 101 and 106. However, the mean squared Z-score weakly correlated with residue distance (*r* = –0.13, *p* = 0.21 for 1H, *r* = –0.34, *p* = 0.003 for 3H, Fig. S6).

We next looked for evidence of “sign epistasis,” where one mutation reverses the sign of the effect of another mutation (Fig. 2C). We defined a Z-score for a single mutation A quantifying the beneficial effect of that mutation relative to the noise, *Z_A_* = (*F*_WT_ — *F_A_*)*σ_A_*, where *F*_WT_ and *F_A_* are the wildtype and mutant free energies, and *σ* is the measurement error estimated as before. Since we are only interested in the sign of the effect, we keep single mutants at the reliable readout boundary. An equivalent Z-score was defined for a mutation A in the background of an existing mutation B: 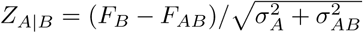, where *F_AB_* is the free energy of the double mutant AB. Significant sign epistasis was defined by *Z*_*A*|*B*_*Z_A_* < 0 and |*Z*_*A*|*B*_|, |*Z_A_*| > 1.64, and reciprocal sign epistasis by the additional symmetric condition *A* ↔ *B*.

We found 44 cases of significant epistasis, listed in `S1_table_sign_epistasis.csv` and summarized in Tables S1 and S2. Deleterious sign epistasis was exceptional, with just one instance in 1H and 4 in 3H (Fig. S4). The four most significant cases of beneficial sign epistatis for each domain are depicted in Fig. 2D. Among cases where both single mutations were deleterious, we found 3% of mutants in 1H and 0.7% in 3H with significant beneficial epistasis, versus 0.06% expected by chance (the null expectation, which takes into account the constraint that *Z_A_*+*Z*_*B*|*A*_ = *Z_B_*+*Z*_*A*|*B*_, is defined in the Methods); 0.7% were reciprocal in 1H, and 0.3% in 3H, versus 0.01% expected by chance. To evaluate how these epistatic interactions may affect affinity maturation, we estimated how often “viable” double mutants were separated from the wildtype by nonviable single mutants, where viability is defined by *K_D_* < 10^−6^M [43-45], forming possible roadblocks to affinity maturation. This strong instance of “rescue” epistasis occured in roughly half of the mutants with beneficial sign epistasis (Table S1 and Table S2).

### Modeling epistasis and its impact on affinity maturation

To integrate the observed epistatic interactions into a predictive model of affinity, we introduced a model of binding free energy as:

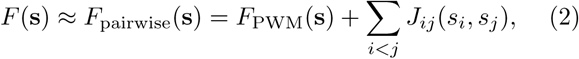

where *J* is the interaction strength between residues. To avoid overfitting and allow for independent validation (in the absence of a sufficient number of triple mutants), we grouped residues into 4 biochemical categories [46] (polar, nonpolar, acidic, basic, see Methods) and let the entries of *J* only depend on that category.

We trained the model on the 1208 1H or 1216 3H double and triple mutants, using a Lasso penalty to control for overfitting. The optimal penalty was set by 10 fold cross-validation, i.e. by maximizing the explained variance of a subset comprising 1/10 of the mutants by using a model trained on the remaining 9/10, averaged over the 10 subsets (Fig. S7A and Methods). Interacting pairs with posterior probabilities > 0.95 as determined by Bayesian Lasso [47] are shown in Fig. 3A and Fig. 3B.

**Fig. 3.**
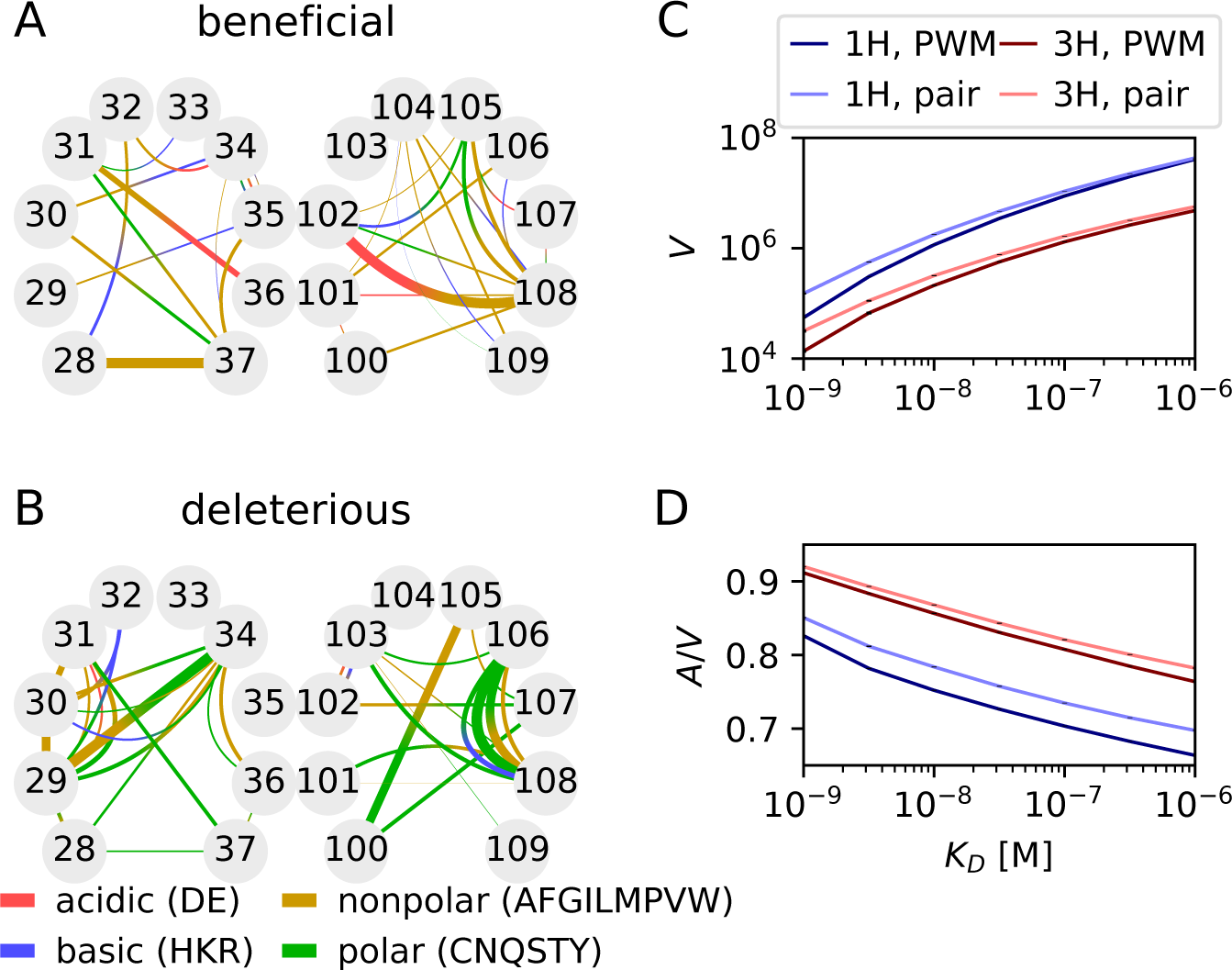
Coarse-grained epistatic model. A model of biochemical epistatic interactions between polar, nonpolar, acidic, and basic residues was fitted to the data using LASSO regularization and tested by cross-validation (Fig. S7A), yielding 1058 CDR1H and 1066 CDR3H interaction terms. Mean **(A)** beneficial and **(B)** deleterious interactions calculated by averaging over all double mutants, colored by interaction type. Line width denotes interaction strength. The model performance on significantly epistatic pairs of positions is shown in Figs. S7B-C, and the number of non vanishing parameters as a function of the significance threshold on the posterior is shown in Fig. S7D. **(C)** Number *V* of amino-acid sequences of the 1H (blue) and 3H (red) regions with dissociation constant below *K_D_*, as estimated by the PWM model (dark color) or the epistatic model (light color). Epistasis enlarges the number of variants with good affinity for both 1H and 3H. **(D)** Mutational flux *A* (defined as the average number of random mutation events from all possible sequences to cause the dissociation constant to cross *K_D_*), normalized by *V*, showing that epistasis also increases the accessibility of the region of good binders in sequence space. Differences between the PWM and epistatic models were robust to errors in the estimate of the interaction parameters (*p* < 10^−5^, Jackknife analysis).

Out of the 720 possible terms, 52 1H and 45 3H interaction terms were identified by this method. Although these interactions, whose number is limited by the number of measured variants, only modestly improved the explained variance relative to the PWM in all multiple mutants (from 62% to 64% for 1H and from 58% to 60% for 3H), it substantially improved the affinity prediction of the mutants with significant epistasis (*R*^2^ from 27% to 50% in 1H, from 13% to 44% for 3H, Fig. S7B-Fig. S7C). Notably, two mutations of the super-optimized 4m5.3 antibody are predicted by the model to have epistatic interactions: a slightly deleterious effect between *A*_101_ and *L*_108_, and a strongly beneficial one between *S*_102_ and *L*_108_.

Next we used our models to estimate the diversity, or “degeneracy”, of antibodies with good binding affinity. Specifically, we evaluated the degeneracy volume *V* of high-affinity sequences as the number of sequences with *K_D_* < B, using either the PWM (Eq. 1) or pairwise (Eq. 2) models, using a combination of exhaustive and Monte-Carlo sampling (Methods). Compared to the coarse-grained pairwise model trained previously, the interaction strength *J* was learned directly for each residue pair, without grouping by biochemical category and with no Lasso penalty. The volume of 1H mutants was larger than that of 3H mutants (Fig. 3C), in agreement with the fact that CDR3H plays a more important role in binding affinity. Epistasis increased the recognition volume for both domains, consistent with the previous observation that epistatic effects are, on average, more beneficial than deleterious. To explore the diversity of evolutionary paths leading to recognition, we computed the mutational flux *A* in and out of the high-affinity region as the probability that a random mutation in a high-affinity sequence (*K_D_* < *B*) causes loss of recognition (*K_D_* > *B*), summed over all high-affinity sequences (Methods). Again we found that epistasis increased the mutational flux, even after normalizing by volume, *A/V* (Fig. 3D). We checked that these differences were robust to sampling noise and overfitting by performing a jack- knife analysis (*p* < 10^−5^ for the difference in *A* and *V* between the PWM and pairwise models, see Methods).

## DISCUSSION

By analyzing massively parallel affinity measurements obtained by Tite-Seq, we painted a detailed picture of epistasis in a well-defined physical phenotype — the binding free energy of an antibody to an antigen. We showed that antibody sequences contain many epistatic interactions contributing to the binding energy, and that many of them have beneficial effects. Our approach involves first training an additive (PWM) model as a baseline, and identifying departures from that model as epistasis. In this comparison, a crucial step was to correct for the two issues of scale and measurement noise.

The first issue, identified by Fisher [36] and also called unidimensional epistasis [25], is the idea that an epistatic trait becomes additive upon a different parametrization. For instance, protein stability, which often determines fitness, is a nonlinear function of the folding free energy difference, which is expected to be roughly additive [12, 27-29, 48-50]. This leads to both a law of diminishing returns [29] and robustness to mutations when the protein is very stable [48]. To disentangle these potential artifacts, we defined our PWM on the binding free energy, which we expect to be additive in sequence content, and we checked that this parametrization was close to minimizing epistasis.

To tackle the second and perhaps more important issue of noise, especially in the context of deep mutational scans where many variants are tested [31], we developed a robust methodology based on Z-scores to identify epistatic interactions as significant outliers. This analysis showed that almost all of the variance unexplained by additivity (∽ 40%) could be attributed to epistasis, making its contribution to the phenotype comparable to that of single mutations. A large fraction of that epistasis was beneficial, in contrast with previous reports of mostly negative epistasis owing to the concavity of the scale [24, 29, 49], which we here circumvent by directly considering the free energy.

Epistasis is key to understanding the predictability and reproducibility of evolutionary paths [18, 51]. Our results show how it could constrain the space of possible hypermutation trajectories during affinity maturation, with important consequences for antibody and vaccine design, as the importance of eliciting responses of antibodies that are not just strongly binding but also evolvable is being increasingly recognized [52]. Targeting epistatic interactions may provide an alternative strategy for optimizing antibody affinity: among the 12 epistatic hotspots in CDR1H and 20 in CDR3H that we identified 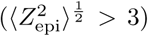, 4 involved positions mutated in the super-optimized 4m5.3 antibody sequence, with a higher epistatic contribution than expected by chance. We also identified 3 cases of beneficial sign epistasis, in which the double mutant was fit despite the single mutant being deleterious. For instance, the D108E mutations in 4m5.3 is deleterious by itself but is rescued beyond the wildtype value by the S101A mutation [19], which occurred first in the directed evolution process [37]. We report 10 extreme cases of viable double mutants whose single-mutant intermediates are nonviable, possibly blocking affinity maturation. However, our analysis of the volume and mutational flux of the region of low binding free energies in sequence space suggests that epistasis facilitates the evolution of high-affinity antibodies. Additionally interactions with the non-mutated parts of the sequence and evolution of the antigen binding partner can either add further constraints or open up additional paths.

Taken together, our results show the importance of taking into account epistasis when predicting antibody evolution and guiding vaccine design. Our systematic approach for identifying and quantifying epistasis, with the implementation of important controls for scale and noise, could be used by other investigators to analyze deep-mutational scans of protein function.

## METHODS

Values of *K_D_* as measured by Tite-Seq for variants of the 4-4-20 fluorescein-binding antibody [34] can be found at https://github.com/jbkinney/16_titeseq. The scripts used for the analyses presented here are available at https://github.com/rhys-m-adams/epistasis_4_4_20.

### Position Weight Matrix

The amino-acid sequence of the 10 amino acid stretches of the CDR1H or CDR3H domains are denoted by s = (s_1_,…,s_10_). The corresponding 30-long nucleotide sequences are denoted by v. The binding free energy *F*(s) of an amino-acid variant is obtained as the mean over 3 replicate experiments, and over all its synonymous variants:

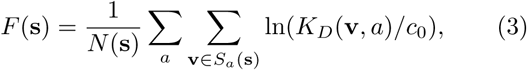

where *S_a_*(s) is the set of measured nucleotide sequences that translate to s in replicate *a*, and *N*(s) = Σ_*a*_|*S_a_*(s)| a normalization constant.

The elements of the PWM are defined as *h_i_*(*q*) = *F*(*s*^(*i,q*)^) − *F_WT_*, where *s*^(*i,q*)^ is the single mutant mutated at position *i* to residue *q*, and *h_i_*(*q*) = 0 when *q* is the wildtype residue at position *i*.

### Optimal nonlinear transformation of the free energy

To test whether transforming *F* through a nonlinear function *E*(*F*) before learning the PWM could improve its predictive power, we defined the nonlinear additive model:

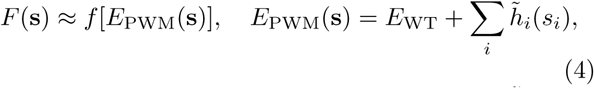

where *f* = *E*^−1^ is the inverse function of *E*, 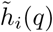 = *E*(s^(*i,q*)^) – *E*_WT_, and *E*(s) is evaluated similarly to Eq. 3: *E*(s) = (1/*N*(s)) Σ_*a*_Σ_v∊*S*_*a*_(s)_ *E*[ln(*K_D_*(v)/*c*_0_)].

To find the transformation *E* that gives the highest explained variance while avoiding overfitting, we aimed to minimize the following objective function:

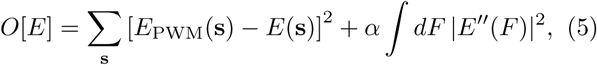

where the sum in s runs over double and triple mutants, and *α* is a tunable parameter.

Numerically, we parametrize the function *E*(*F*) as piecewise linear: *E*(*F*) = *E_i_* × (*F*_*i*+1_ – *F*)/*δF* + *E*_*i*+1_ × (*F* – *F_i_*)/*δF* for *F_i_* ≤ *F* ≤ *F_i_*, where *F_i_* are equally spaced grid point along *F*, *δF* = *F*_*i*+1_ – *F_i_*, and *E_i_* the value of *E* at these points. The smoothing penalty is approximated by a sum over the squared discretized second derivative: 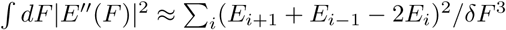.

We minimize *O*[*E*] ≈ *O*[*E*_1_,…, *E_N_*] as a quadratic function of its arguments (*E_i_*), while imposing boundary constraints on the PWM prediction and the requirement that *E* is a increasing function of *F* (i.e. *E*_*i*+1_ > *E_i_*), using the python package `cvxopt` [53].

The hyper-parameter *α* is evaluated by maximizing the generalized cross-validation of the coefficient of determination

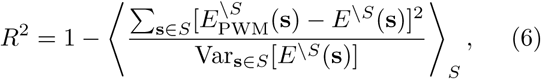

where 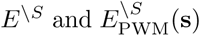 are learned through optimizing Eq. 5, but after removing from the dataset a subset *S* of the multiple mutants comprising one tenth of the total. The average is over ten non-overlapping subsets *S*.

This method was first tested on simulated data. Each PWM element 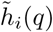 was drawn from a normal distribution of zero mean and variance 1, and then *E*_PWM_(S) was computed for each of the antibody sequences present in our data. Our simulated “measurement” was defined as a function of a noisy realization of *E* = *E*_PWM_ + ∊ (where ∊ is some Gaussian noise) in three different ways: linear *F* = *E*, exponential *F* = exp(*E*), high-frequency *F* = 2*E* + sin(2*E*), and logistic *F* = 1/[1 + exp(−*E*)]. ∊ was drawn from a centered normal distribution with 1/2 the standard deviation of *E*_PWM_. *F* was then truncated to the 200th lowest and 200th highest values, to mimick the boundary cutoff in our measurements. Comparing our original *E*_PWM_ to our fit *Ê* shows that our method is able to infer the true PWM model and a smooth nonlinearity from noisy data (Fig. S2).

We then applied the method to the experimental data. The cross-validation *R*^2^ is represented as a function of the smoothing parameter *α* in Fig. S3A, and the corresponding optimal function *E*(*F*) in Fig. S3B. The comparison between measurement and the PWM model is shown in Fig. S3C.

### Z-scores

We used synonymous mutants to estimate our measurement error. The mean free energy of a nucleotide sequence is defined as the mean over replicate measurements: *F*(v) = 〈ln(*K_D_*(v,a))_*a*_, and the standard error σ(v) is defined accordingly as the pooled error over replicates. Antibodies with *K_D_* having median values at the boundary values of 10^−9.5^ or 10^−5^ were excluded from the analysis since these values artificially cluster at the boundary, leading to underestimates of error.

The error Z-score was calculated between pairs of nucleotide sequences with the same amino acid translation: 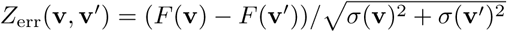.

Epistatic Z-scores were estimated by calculating the measurement error over both replicates and synonymous variants, as in Eq. 3:

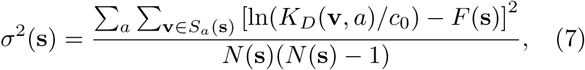

and the pooled standard error for a PWM prediction, calculated as the sum of measurement errors from single mutations:

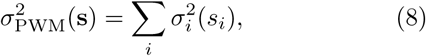

where *σ_i_*(*q*) = *σ*(s^(*i,q*)^), and *σ_i_*(*q*) = 0 when *q* is the wildtype residue at *i*. The epistatic Z-score is defined as:

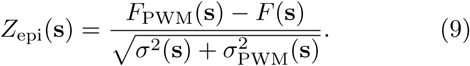

### Null model for sign epistasis

To calculate p-values for sign epistasis, we used the following null model for sets of four Z-scores satisfying *Z_A_* + *Z*_*B*|*A*_ = *Z_B_* + *Z*_*A*|*B*_. Calling *x*_1_ = *Z_A_*, *x*_2_ = *Z*_*B*|*A*_, *x*^3^ = – *Z*_*A*|*B*_, *x*_4_ = – *Z_A_*, the condition becomes that each *x_i_* has zero mean and variance one, with the constraint 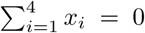. The distribution with maximum entropy satisfying these requirements is a centered multi-variate Gaussian uniquely defined by its covariance matrix 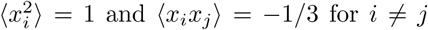. The p-value for sign epistasis, *Z_A_* > 1.64 and *Z*_*A*|*B*_ < 1.64, was estimated by Monte Carlo sampling under that Gaussian distribution as Pr(*x*_1_ > 1.64 & *x*_2_ > 1.64) + Pr(*x*_3_ > 1.64 & *x*_4_ > 1.64) − Pr(*x*_1_ > 1.64 & *x*_2_ < −1.64 & *x*_3_ > 1.64 & *x*_4_ < –1.64) = 6.2 · 10^−4^, and the probability for reciprocal sign epistasis as Pr(*x*_1_ > 1.64 & *x*_2_ < – 1.64 & *x*_3_ > 1.64 & *x*_4_ < –1.64) = 10^−4^.

### Epistatic model

The epistatic terms of the pairwise model were made to depend on the biochemical categories of the interacting residues, 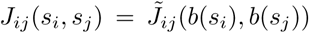, with *b*(*s*) = nonpolar for *s* = AFGILMPVW, *b*(*s*) = polar for *s* = CNQSTY, *b*(*s*) = acidic for *s* = DE, and *b*(*s*) = basic for *s* = HKR. A fifth category was added to correspond to the wildtype residue, so that 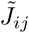 (wildtype, *b*) = 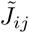 The model was trained by minimizing the mean squared error with a regularization penalty over all matrices 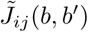:

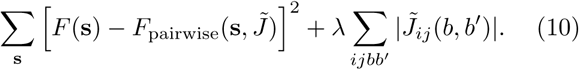

The Lasso penalty λ was learned by 10-fold crossvalidation, and energy terms found in less than 2 sequences were excluded from the fit. Posterior values for 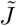 terms were calculated using Bayesian Lasso [47].

The volume and mutational flux were defined as:

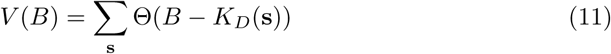

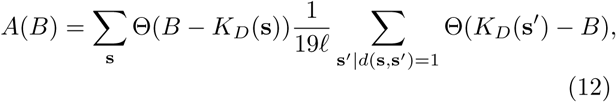

where Θ(*x*) is the Heaviside function, i.e. Θ(*x*) = 1 if *x* ≥ 0 and 0 otherwise; *d*(*s*, *s′*) is the Hamming distance between two sequences; and *l* = 10 is the sequence length. The normalization 19 × *l* corresponds to the number of mutants *s′* at Hamming distance 1 from s. The sums over s in Eqs. 11-12 have 20^10^ elements and are computationally intractable. To overcome this, we approximated the sum using a mixture of Monte-Carlo and complete enumeration, depending on the distance of s from the wildtype. Calling *C_d_* the set of sequences s at Hamming distance *d* from wildtype, we used:

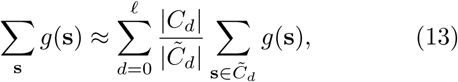

where *g*(s) is a function of s to be summed such as in *V* or *A* in Eqs. 11-12, and 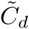 is a random subset of *C_d_* of size min(|*C_d_*|, *P_d_*), with 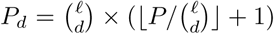, where *P* is the maximum number of sequences one is willing to sample completely aa each d to perform the estimation, and where 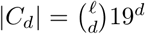. For small *d*, when |*C_d_*| ≤ *P_d_*, the enumeration is complete, while for large *d* and |*C_d_*| > *P_d_*, the sum is estimated from a uniformly distributed Monte Carlo sample of *C_d_*.

## ACKNOWLEDGMENTS

We would like to thank Yuanzhe Guan and Carlos Ta- laveira for their suggestions. The authors declare no conflicts of interest. R.M.A., T.M. and A.M.W. were supported by grant ERCStG n. 306312.

---

**Table S1.** Summary of CDR1H sign epistasis with double mutants. Obs/exp denotes observed sign epistasis events versus expected epistasis events in a Gaussian noise model. Expected numbers below 0.01 are rounded up to 0.01. We measured 1058 CDR1H double mutants in total.

| domain | # of nonviable single mutations | epistasis type | # candidate mutants | # of mutants with sign epistasis (obs/exp) | # of mutants with reciprocal sign epistasis (obs/exp) | # of viable mutants with sign epistasis (obs/exp) | # of viable mutants with reciprocal sign epistasis (obs/exp) | # of sign epistatic mutants with $K_D < \text{WT}$ (obs/exp) |
| --- | --- | --- | --- | --- | --- | --- | --- | --- |
| 1H | 0 | beneficial | 293 | 2/0.21 | 0/0.04 | 2/0.06 | 0/0.01 | 0/0.01 |
| 1H | 1 | beneficial | 532 | 16/0.32 | 0/0.05 | 16/0.09 | 0/0.02 | 2/0.01 |
| 1H | 0, 1 | beneficial | 825 | 18/0.53 | 0/0.09 | 18/0.15 | 0/0.03 | 2/0.02 |
| 1H | 2 | beneficial | 172 | 14/0.10 | 7/0.02 | 12/0.03 | 6/0.01 | 0/0.01 |
| 1H | 0 – 2 | beneficial | 997 | 32/0.63 | 7/0.11 | 30/0.18 | 6/0.03 | 2/0.02 |
| 1H | 0 – 2 | deleterious | 53 | 1/0.63 | 0/0.11 | 0/0.18 | 0/0.03 | 0/0.02 |

**Table S2.**
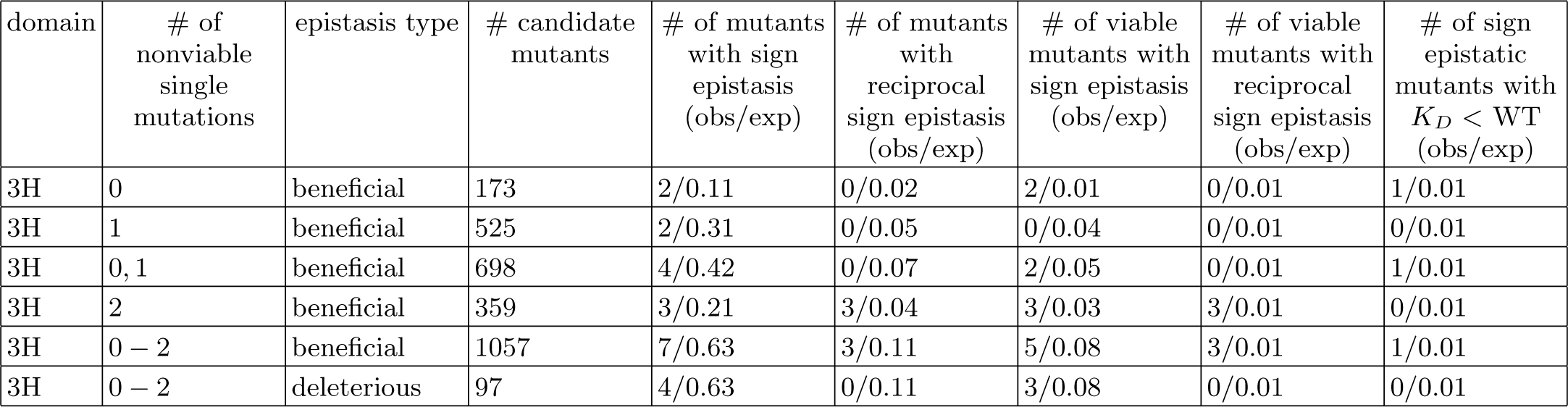
Summary of CDR3H sign epistasis with double mutants. Obs/exp denotes observed sign epistasis events versus expected epistasis events in a Gaussian noise model. Expected numbers below 0.01 are rounded up to 0.01. We measured 1066 CDR3H double mutants in total.

**Fig. S1.**
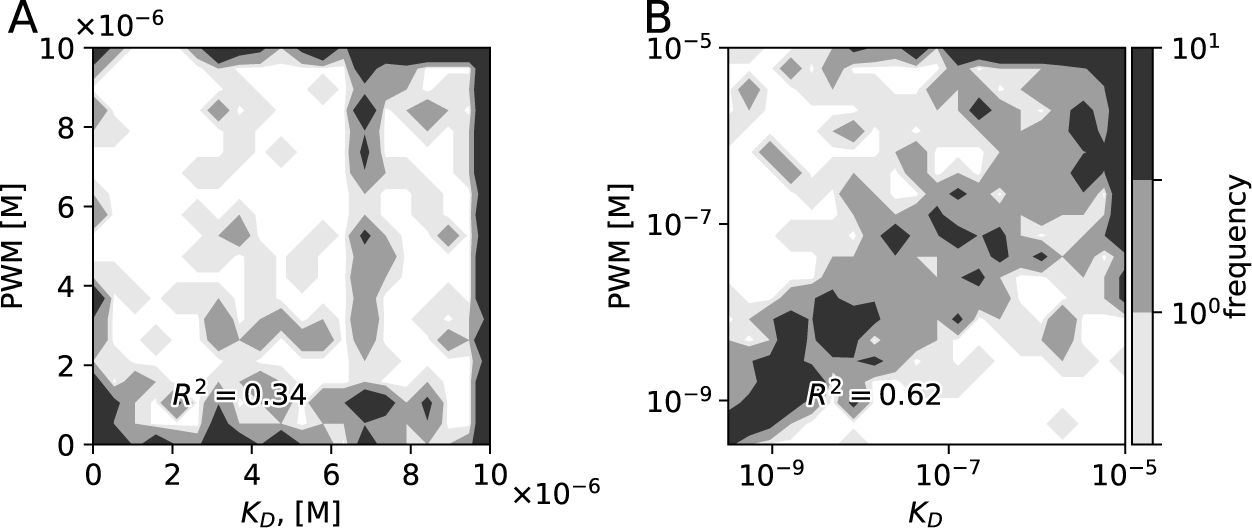
Comparison between data and model prediction of the binding affinity of multiple mutants using PWMs constructed from **A)** *K_D_* and **B)** *F* = ln(*K_D_=c_0_*). *R^2^* denotes the coefficient determination (fraction of explained variance). Although both predictions are very significant (*p* < 10^−5^, F-test), the PWM based on *F* is much better.

**Fig. S2.**
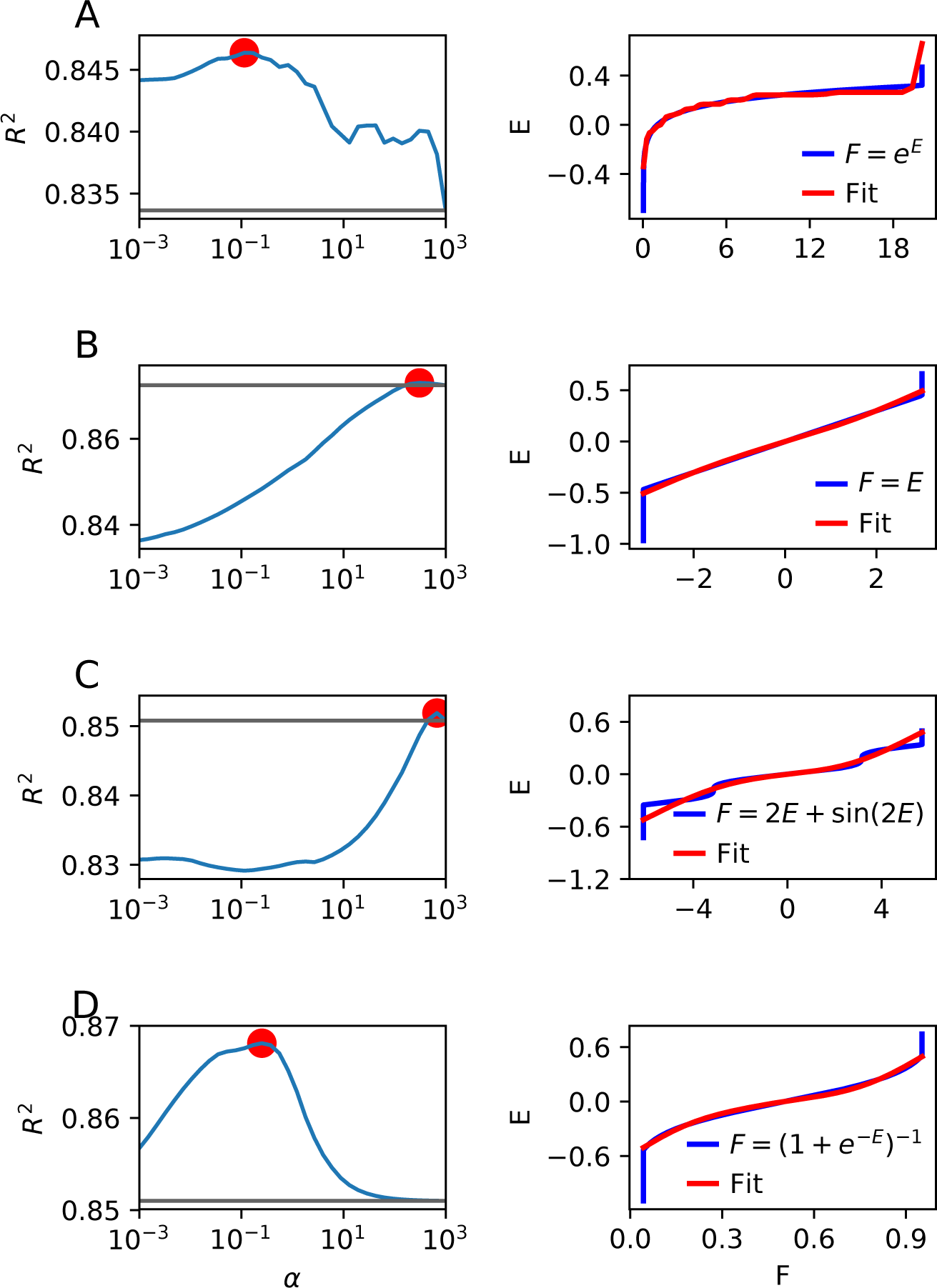
Test of the inference method for learning the nonlinear scale on synthetic data. A PWM was generated randomlly, with terms drawn from a normal distribution, and applied to the sequences from our dataset to simulate a “true” score (*E*_PWM_). Gaussian noise with 50% of the standard deviation of *E*_PWM_ was added. The score was the converted into a free energy *F* = *f*(*E*) (Methods), where *f*, the inverse function of E, was a **(A)** linear, **(B)** exponential, **(C)** high frequency, or **(D)** logistic transformation. The range of measured *F* was cut off at the boundaries as in the data, resulting in the vertical lines in the right panels. Left panels: cross-validated fraction of explained variance, as a function of the regularization parameter *α* penalizing the second derivative of the function *E*. Large *α* implies very smooth functions, while small *α* allows for high-frequency variations. These results the original PWM can be recovered as well as the non-linearity, except for its high-frequency components (C).

**Fig. S3.**
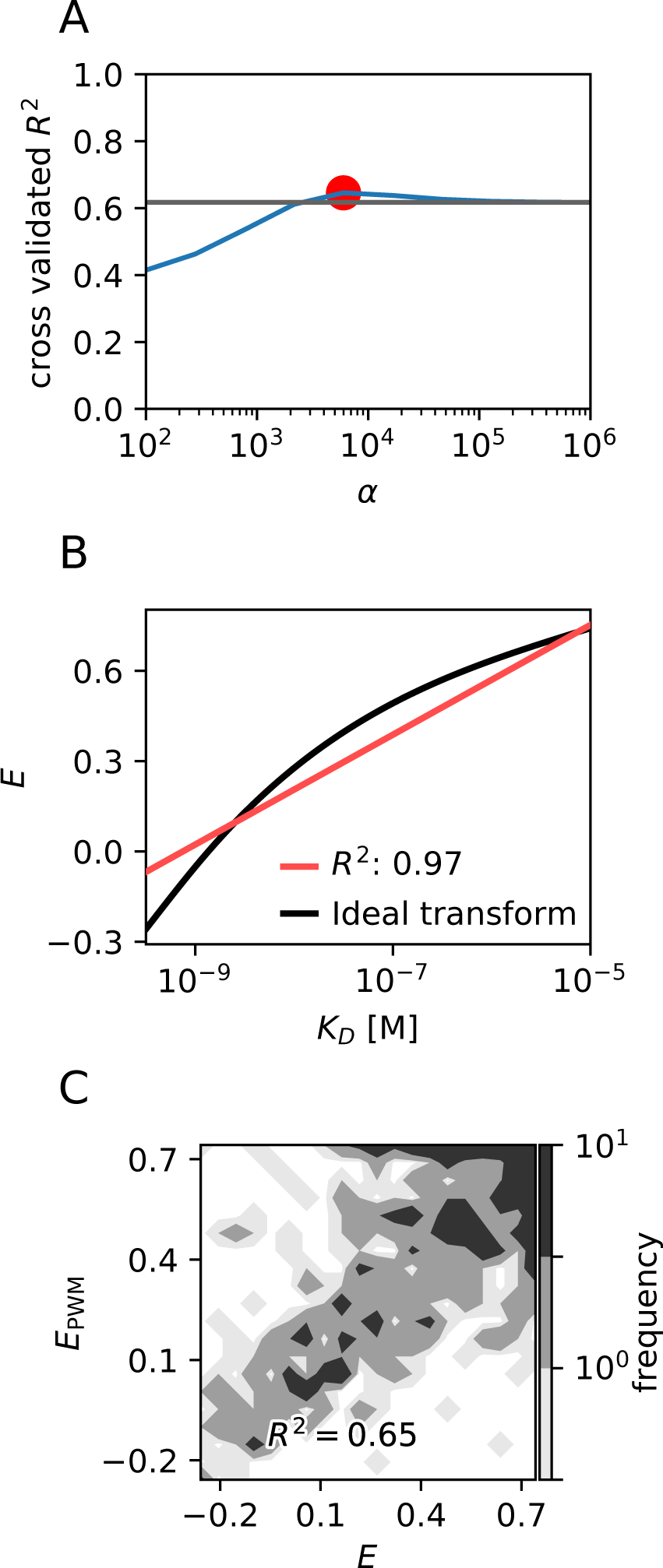
Optimizing the non-linear scale *E* for the PWM model on real data. **(A)** Cross-validated fraction of explained variance, as a function of the regularization parameter *α* penalizing the second derivative of the function *E*. **(B)** Optimized non-linear scale *E* as a function of *F* = ln(*K_D_/c*_0_) (black), compared to identity (red). **(C)** Comparison between data and the PWM model with the optimal nonlinear scale.

**Fig. S4.**
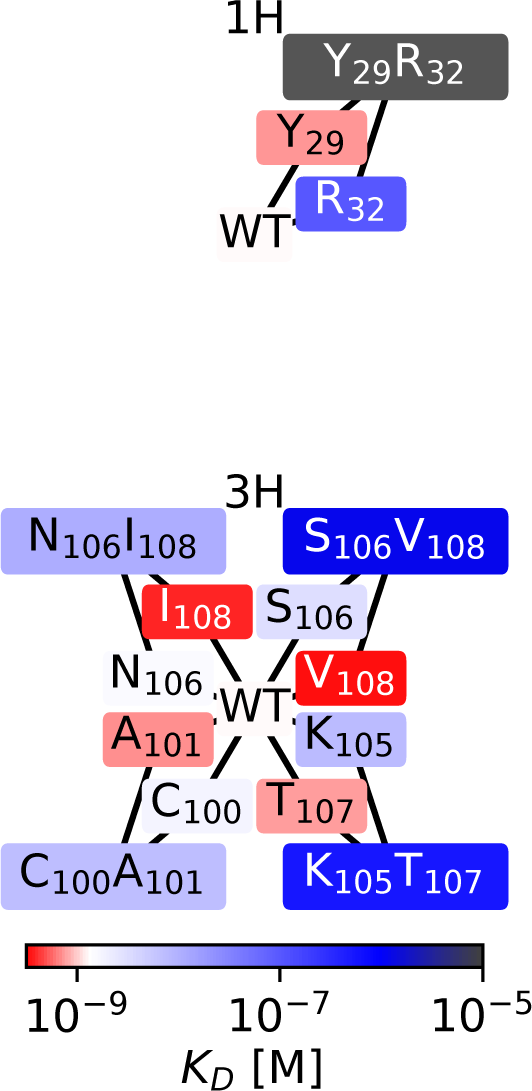
All deleterious sign epistasis examples are shown for the 1H and 3H domains.

**Fig. S5.**
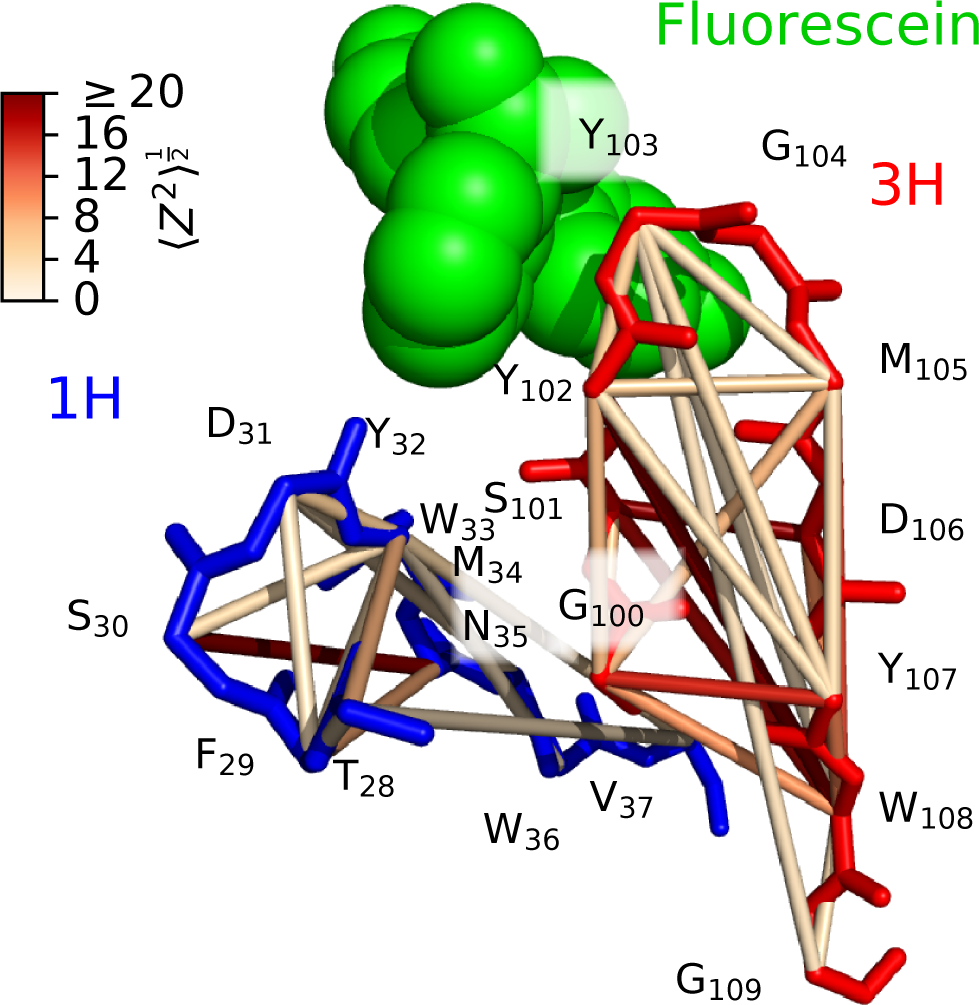
Pairs of positions with standard epistatic effect 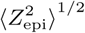 > 3 (with the mean taken over all measured amino-acid variants at the two positions) are superimposed on the wildtype antibody structure.

**Fig. S6.**
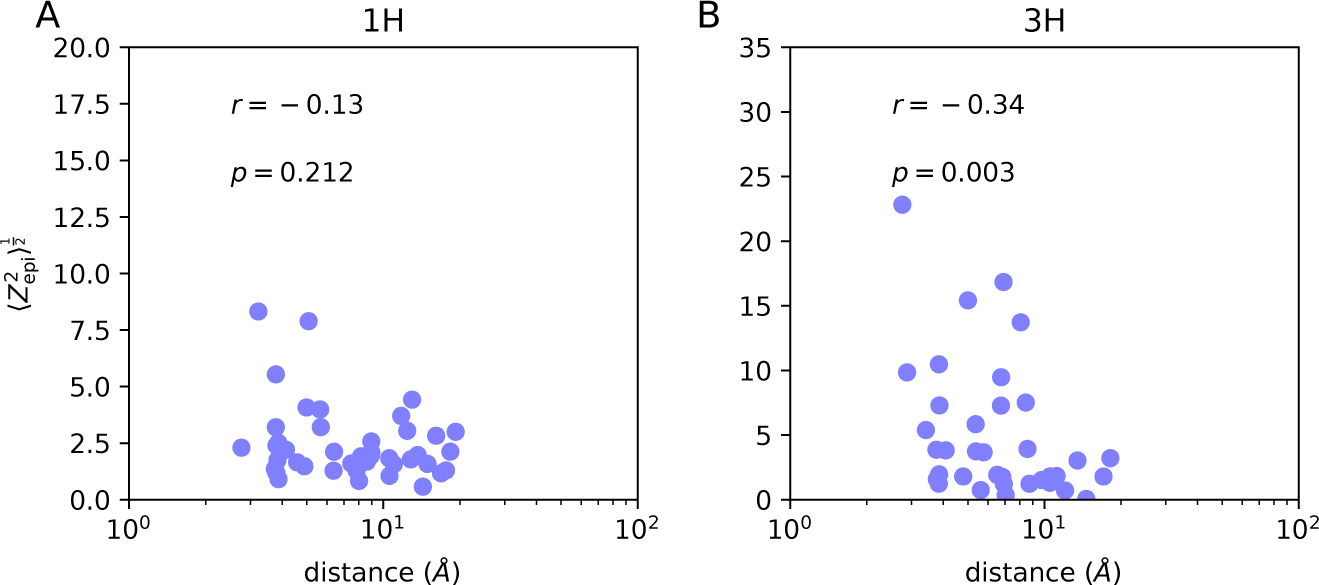
Standard epistatic effect 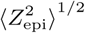 versus distance between residues in the **(A)** 1H and **(B)** 3H domains.

**Fig. S7.**
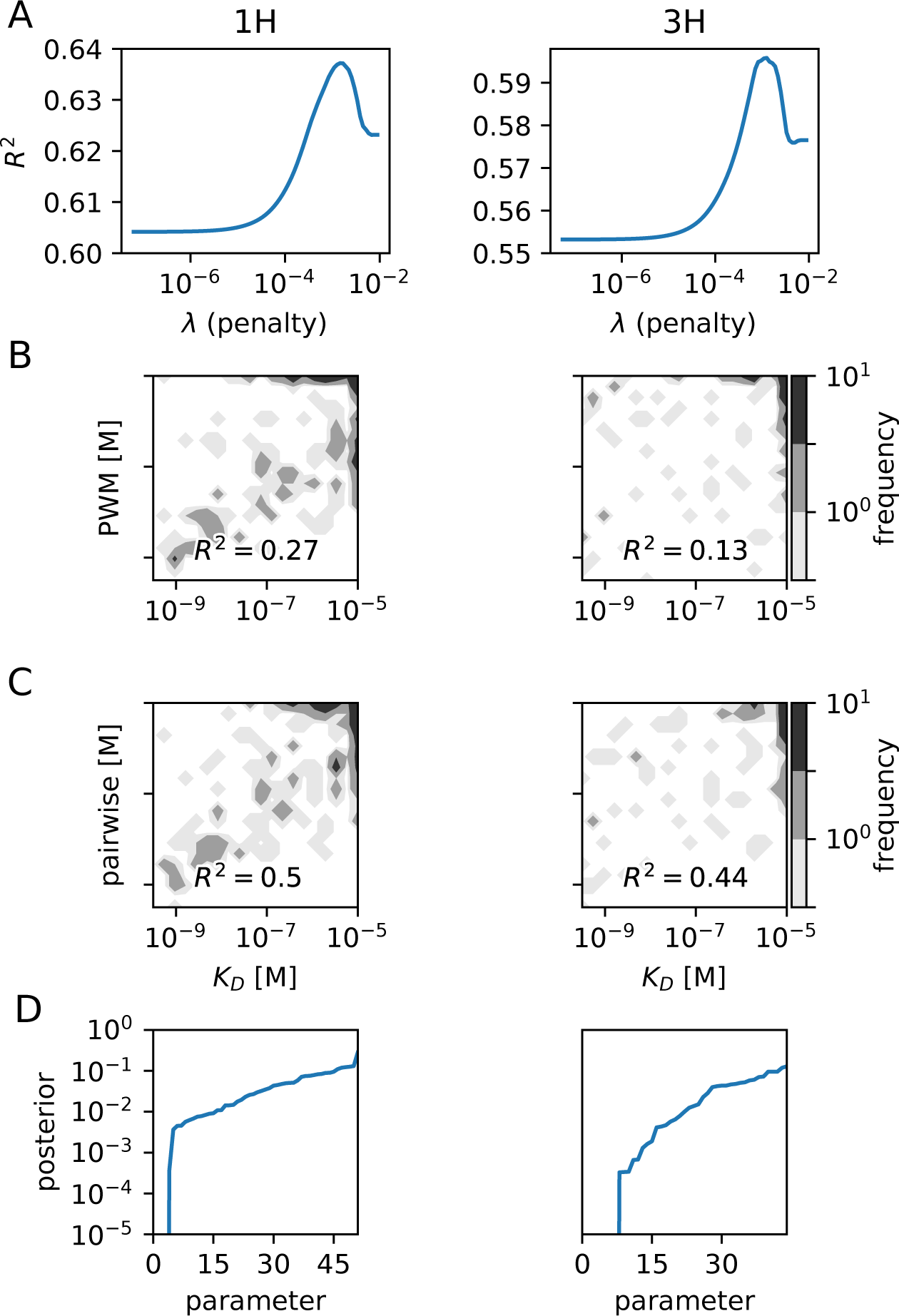
Inferrence of the epistatic model. Left panels correspond to the 1H domain, and right panels to the 3H domain. Parameters were fit by minimizing the mean squared model error with a L1 penalty on the parameters with coeffcient λ. **(A)** The cross-validated coeffcient of determination (1 – standard error^2^/standard deviation^2^) has a clear maximum as a function of λ. **(B)** and **(C)** Model prediction versus measurement of *F* = ln(*K_D_=c_0_*), for sequences involving a non-zero interacting term, when using **(B)** the PWM model and **(C)** the epistatic model with optimal λ. **(D)** Rank-ordered posterior probability of 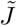 parameters to be non-zero, according to the Bayesian Lasso posterior [47]. Monte Carlo Markov Chains (MCMC) were used to estimate posterior probability of a parameter switching signs. Multiple (2 × number of non-zero parameters from Lasso optimization + 2) MCMCs were initialized (“thermalized”) for 227 steps × the number of parameters. After thermalization, the MCMCs were simulated for another 454 steps × the number of parameters. Parameter values were sampled at intervals where their autocorrelations were ≤ 0.1. The posterior probabilty was calculated as the fraction of time the parameter switched signs in the MCMCs.

## References

[1] Cobey S, Wilson P, Iv FAM, Cobey S (2015) The evolution within us.

[2] Eisen HN, Siskind GW (1964) Variations in A_nities of Antibodies during the Immune Response. Biochemistry 3:996–1008.

[3] Corti D, Lanzavecchia A (2013) Broadly Neutralizing Antiviral Antibodies Vol. 31, pp 705–742.

[4] Esmaielbeiki R, Krawczyk K, Knapp B, Nebel JC, Deane CM (2016) Progress and challenges in predicting protein interfaces. Brief. Bioinform. 17:117–131.

[5] Phillips PC (2008) Epistasis - The essential role of gene interactions in the structure and evolution of genetic systems. Nat. Rev. Genet. 9:855–867.

[6] Carter AJR, Hermisson J, Hansen TF (2005) The role of epistatic gene interactions in the response to selection and the evolution of evolvability. Theoretical Population Biology 68:179–196.

[7] Good BH, Desai MM (2015) The impact of macroscopic epistasis on long-term evolutionary dynamics. Genetics 199:177–190.

[8] Paixão T, Barton NH (2016) The e_ect of gene interactions on the long-term response to selection. Proc. Natl. Acad. Sci. 113:4422–4427.

[9] Breen MS, Kemena C, Vlasov PK, Notredame C, Kondrashov Fa (2012) Epistasis as the primary factor in molecular evolution. Nature 490:535–538.

[10] Weinreich DM, Delaney NF, DePristo MA, Hartl DL (2006) Darwinian evolution can follow only very few mutational paths to fitter proteins. Science 312:111–114.

[11] Poelwijk FJ, Kiviet DJ, Weinreich DM, Tans SJ (2007) Empirical fitness landscapes reveal accessible evolution evolutionary paths. Nature 445:383–386.

[12] Gong LI, Suchard MA, Bloom JD (2013) Stability-mediated epistasis constrains the evolution of an influenza protein. Elife 2013:1–19.

[13] Anderson DW, McKeown AN, Thornton JW (2015) Intermolecular epistasis shaped the function and evolution of an ancient transcription factor and its DNA binding sites. Elife 4:1–26.

[14] Podgornaia AI, Laub MT (2015) Protein evolution. pervasive degeneracy and epistasis in a protein-protein interface. Science 347:673–677.

[15] Bloom JD, Labthavikul ST, Otey CR, Arnold FH (2006) Protein stability promotes evolvability. Proc. Natl. Acad. Sci. 103:5869–5874.

[16] Bloom JD, Gong LI, Baltimore D (2010) Permissive Secondary Mutations Enable the Evolution of Influenza Oseltamivir Resistance. Science (80-.). 328:1272–1275.

[17] Chou HH, Chiu HC, Delaney NF, Segrè D, Marx CJ (2011) Diminishing returns epistasis among beneficial mutations decelerates adaptation. Science 332:1190–1192.

[18] Kryazhimskiy S, Rice DP, Jerison ER, Desai MM (2014) Global epistasis makes adaptation predictable despite sequence-level stochasticity. Science (New York, N.Y.) 344:1519–1522.

[19] Midelfort KS, et al. (2004) Substantial energetic improvement with minimal structural perturbation in a high affinity mutant antibody. J. Mol. Biol. 343:685–701.

[20] Koenig P, et al. (2015) Deep sequencing-guided design of a high affinity dual specificity antibody to target two angiogenic factors in neovascular age-related macular degeneration. Journal of Biological Chemistry 290:21773–21786.

[21] Boyer S, et al. (2016) Hierarchy and extremes in selections from pools of randomized proteins. Proc. Natl. Acad. Sci. 113:3482–3487.

[22] Mora T, Walczak AM, Bialek W, Callan CG (2010) Maximum entropy models for antibody diversity. Proc. Natl. Acad. Sci. 107:5405–5410.

[23] Asti L, Uguzzoni G, Marcatili P, Pagnani A (2016) Maximum-Entropy Models of Sequenced Immune Repertoires Predict Antigen-Antibody Affinity. PLOS Comput. Biol. 12:e1004870.

[24] Schenk MF, et al. (2013) Patterns of epistasis between beneficial mutations in an antibiotic resistance gene. Mol. Biol. Evol. 30:1779–1787.

[25] Szendro IG, Schenk MF, Franke J, Krug J, De Visser JAGM (2013) Quantitative analyses of empirical fitness landscapes. J. Stat. Mech. Theory Exp. 2013.

[26] de Visser JAGM, Krug J (2014) Empirical fitness landscapes and the predictability of evolution. Nature Reviews Genetics 15:480–490.

[27] Sarkisyan KS, et al. (2016) Local fitness landscape of the green fluorescent protein. Nature 533:397–401.

[28] Jacquier H, et al. (2013) Capturing the mutational landscape of the beta-lactamase TEM-1. Proc. Natl. Acad. Sci. 110:13067–13072.

[29] Bank C, Hietpas RT, Jensen JD, Bolon DNA (2015) A Systematic Survey of an Intragenic Epistatic Landscape. Mol. Biol. Evol. 32:229–238.

[30] Fowler DM, Fields S (2014) Deep mutational scanning: a new style of protein science. Nat. Methods 11:801–807.

[31] Araya CL, et al. (2012) A fundamental protein property, thermodynamic stability, revealed solely from large-scale measurements of protein function. Proc Natl Acad Sci USA 109:16858–16863.

[32] Olson CA, Wu N, Sun R (2014) A comprehensive biophysical description of pairwise epistasis throughout an entire protein domain. Current Biology 24:2643–2651.

[33] Vodnik M, Zager U, Strukelj B, Lunder M (2011) Phage display: Selecting straws instead of a needle from a haystack. Molecules 16:790–817.

[34] Adams RM, Mora T, Walczak AM, Kinney JB (2016) Measuring the sequence-affinity landscape of antibodies with massively parallel titration curves. eLife 5:e23156.

[35] Wells JA (1990) Additivity of mutational effects in proteins. Biochemistry 29:8509–8517.

[36] Fisher R (1918) The Correlation between Relatives on the Supposition of Mendelian Inheritance. Trans. R. Soc. Edinburgh 52:399–433.

[37] Boder ET, Midelfort KS, Wittrup KD (2000) Directed evolution of antibody fragments with monovalent femtomolar antigen-binding affinity. Proc. Natl. Acad. Sci. 97:10701–10705.

[38] Romero PA, Krause A, Arnold FH (2013) Navigating the protein fitness landscape with gaussian processes. Proc. Natl. Acad. Sci. 110:E193–E201.

[39] Morcos F, et al. (2011) Direct-coupling analysis of residue coevolution captures native contacts across many protein families. Proc. Natl. Acad. Sci. 108:E1293–E1301.

[40] McLaughlin RN, Poelwijk FJ, Raman A, Gosal WS, Ranganathan R (2012) The spatial architecture of protein function and adaptation. Nature 491:138–142.

[41] Zhang X, Perica T, Teichmann SA (2013) Evolution of protein structures and interactions from the perspective of residue contact networks. Current Opinion in Structural Biology 23:954–963.

[42] Melamed D, Young DL, Gamble CE, Miller CR, Fields S (2013) Deep mutational scanning of an RRM domain of the saccharomyces cerevisiae poly(a)-binding protein. RNA 19:1537–1551.

[43] Batista FD, Neuberger MS (1998) Affinity dependence of the b cell response to antigen: A threshold, a ceiling, and the importance of off-rate. Immunity 8:751–759.

[44] Foote J, Eisen HN (1995) Kinetic and affinity limits on antibodies produced during immune responses. Proc. Natl. Acad. Sci. USA 92:1254–1256.

[45] Roost HP, et al. (1995) Early high-affinity neutralizing anti-viral igg responses without further overall improvements of affinity. Proc. Natl. Acad. Sci. USA 92:1257–1261.

[46] Voet D, Voet JG (2011) Biochemistry, 4th edition. New York: John Wiley & Sons Inc pp 68–69.

[47] Park T, Casella G (2008) The bayesian lasso. Journal of the American Statistical Association 103:681–686.

[48] Bloom JD, et al. (2005) Thermodynamic prediction of protein neutrality. Proc. Natl. Acad. Sci. 102:606–611.

[49] Bershtein S, Segal M, Bekerman R, Tokuriki N, Tawfik DS (2006) Robustness-epistasis link shapes the fitness landscape of a randomly drifting protein. Nature 444:929–932.

[50] Serohijos AW, Shakhnovich EI (2014) Merging molecular mechanism and evolution: Theory and computation at the interface of biophysics and evolutionary population genetics. Curr. Opin. Struct. Biol. 26:84–91.

[51] Lässig M, Mustonen V, Walczak AM (2017) Predicting evolution. Nature Ecology & Evolution 1:0077.

[52] Wang S, et al. (2015) Manipulating the selection forces during affinity maturation to generate cross-reactive HIV antibodies. Cell 160:785–797.

[53] Andersen MS, Dahl J, Vandenberghe L (2013) Cvxopt: A python package for convex optimization, version 1.1. 6. Available at cvxopt. org 54.

